# Isolation and Probiotic Characterization of Lactic Acid Bacteria from the Phylloplanes of Edible and Culinary Leaves

**DOI:** 10.1101/2025.04.17.649317

**Authors:** S Shashank, B V Uday Kumar, Uddalak Das

## Abstract

The phylloplane of edible and culinary plants hosts diverse microbial communities, including underexplored lactic acid bacteria (LAB) with probiotic potential. This study isolated LAB from the leaf surfaces of eight plant species and evaluated their probiotic attributes. Isolates were tested for acid, bile, NaCl, and phenol tolerance; temperature resistance; haemolytic activity; antibiotic susceptibility; auto-aggregation; and hydrophobicity. All showed γ-haemolysis, indicating non-pathogenicity. *Lactobacillus acidophilus* and LAB-10 demonstrated strong acid (pH 2.0–3.5) and bile tolerance, high hydrophobicity (72% and 60%), and strong auto-aggregation (42% and 36%). They tolerated up to 6% NaCl, 0.4% phenol, and grew optimally at 37–40□°C. Antibiotic resistance varied among isolates. These findings reveal the phylloplane as a promising reservoir of functionally potent probiotic LAB strains for potential use in functional foods and therapeutics.

**Graphical abstract:** 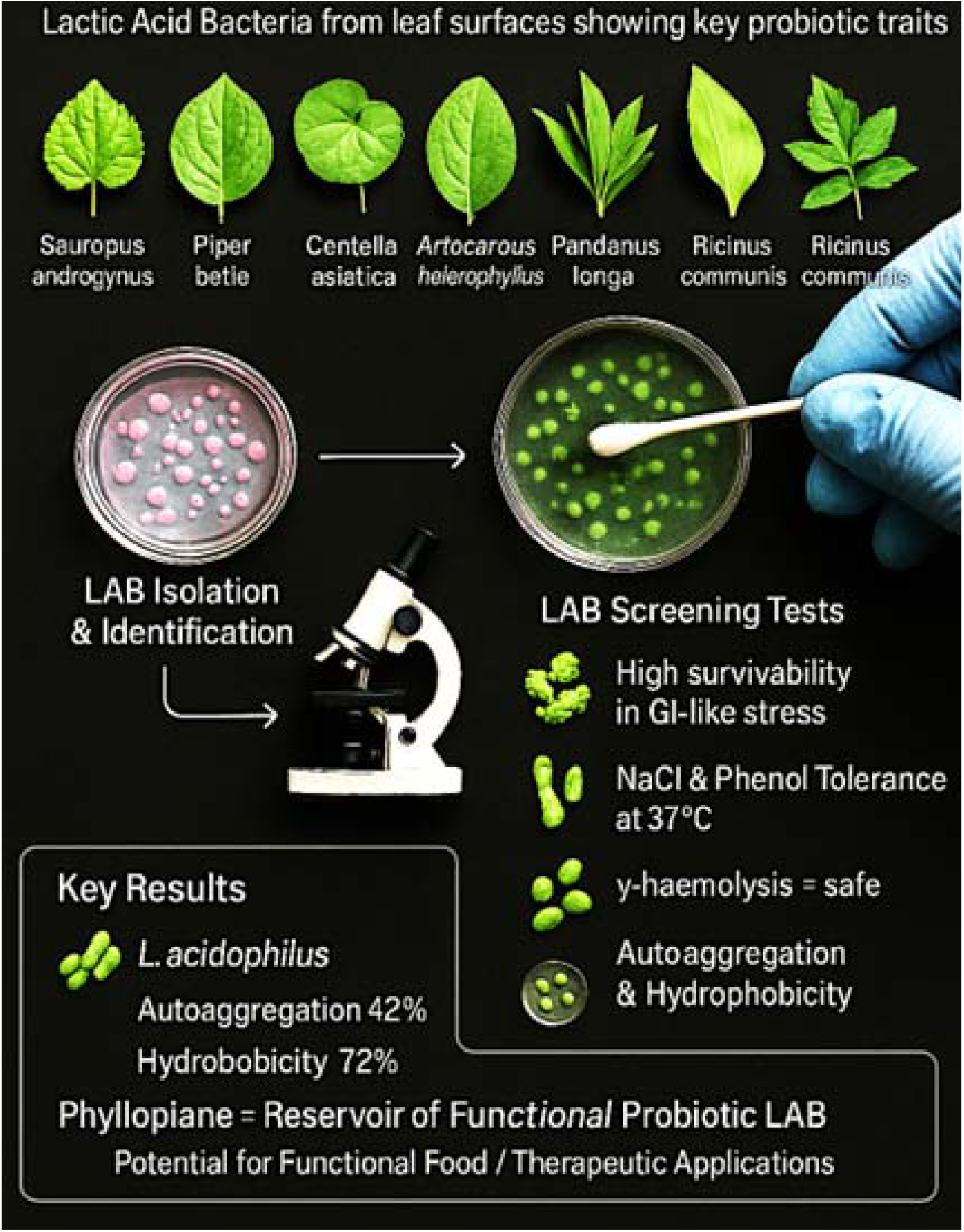

Illustration of the isolation and screening of lactic acid bacteria (LAB) from the phylloplanes of edible and culinary plants, highlighting key probiotic traits. Eight plant species served as microbial reservoirs, with LAB isolates evaluated for gastrointestinal stress tolerance, NaCl and phenol resistance, non-hemolytic (γ-haemolysis) behavior, auto-aggregation, and hydrophobicity. *Lactobacillus acidophilus* exhibited outstanding probiotic potential with 72% hydrophobicity and 42% auto-aggregation. This study reveals the phylloplane as a promising source of functional probiotic LAB with potential applications in functional foods and therapeutic development.

## 1. Introduction

The phylloplane, or aerial surface of plant leaves, provides a diverse and dynamic habitat for microbial colonization, including bacteria, fungi, and viruses. This niche supports a complex microbial community influenced by leaf morphology, age, physicochemical properties, and environmental factors such as humidity and temperature. These factors collectively shape the microbial diversity and functionality on the leaf surface, making it a critical interface between plants and the surrounding environment (Das, Chandramouli, *et al*., 2025; Massoni *et al*., 2021; Remus□Emsermann *et al*., 2014; Vorholt *et al*., 2017). A variety of leafy plants such as *Sauropus androgynus* (Chekkurmanis), *Piper betle* (Betel vine), *Centella asiatica* (Brahmi), *Artocarpus heterophyllus* (Jackfruit), *Pandanus amaryllifolius* (Pandan), *Musa* sp. (Banana), *Curcuma longa* (Turmeric), and *Ricinus communis* (Castor) are commonly consumed raw or used in culinary practices. These leaves are valued not only for their nutritional content, including proteins, essential minerals, and dietary fibre, but also for their bioactive properties such as antioxidant, antimicrobial, anti-inflammatory, and neuroprotective effects (Naik *et al*., 2021; Shanmugapriya *et al*., 2022). Their traditional medicinal use includes lactation enhancement, cognitive support, blood pressure regulation, and improved skin health.

Lactic acid bacteria (LAB) are Gram-positive, non-sporulating, facultative anaerobes that primarily ferment carbohydrates to produce lactic acid. They are ubiquitous in nature and have been isolated from diverse sources including fermented foods, plant surfaces, animal mucosa, and soil (Duar *et al*., 2017; Zheng *et al*., 2020). LAB play a central role in the fermentation of a wide array of food products, enhancing preservation, flavor, and nutritional value (Wuyts *et al*., 2018). Major LAB genera include *Lactobacillus, Lactococcus, Leuconostoc, Pediococcus, Enterococcus*, and *Weissella*, many of which have undergone recent taxonomic reclassification (Zheng *et al*., 2020). LAB associated with raw fruits and vegetables are of particular interest due to their potential as natural probiotics. Probiotics are defined as “live microorganisms which, when administered in adequate amounts, confer a health benefit on the host” (Hill *et al*., 2014). Many strains of *Lactobacillus* and *Bifidobacterium* have demonstrated probiotic properties, including antimicrobial activity, immunomodulation, and restoration of gut microbial balance (Das, Chanda, *et al*., 2024; Plaza-Diaz *et al*., 2020). Their inhibitory effect against intestinal pathogens is largely attributed to the production of lactic acid, hydrogen peroxide, bacteriocins, and other bioactive compounds, as well as competitive exclusion mechanisms (Chugh & Kamal-Eldin, 2020).

For a LAB strain to be considered probiotic, it must demonstrate acid and bile tolerance, adhesion to intestinal epithelial cells, and resistance to digestive enzymes (Das, Uttarkar, *et al*., 2025; Petrova *et al*., 2022). In addition, probiotic strains are known to synthesize B-group vitamins, short-chain fatty acids, and essential amino acids, and may produce enzymes such as esterases and lipases that aid host metabolism (Cizeikiene *et al*., 2020; Riaz Rajoka *et al*., 2017). Several LAB strains have also been reported to produce antimicrobial peptides with potential immunosuppressive or anticancer effects (Das, Banerjee, *et al*., 2024; Umu *et al*., 2020). The GRAS (Generally Recognized as Safe) status granted by the U.S. FDA and the QPS (Qualified Presumption of Safety) status by the EFSA further support their widespread use in food and therapeutic applications.

Given the growing consumption of raw leafy vegetables and their role as carriers of beneficial microbes, the phylloplane represents a promising yet underexplored niche for the isolation of LAB with probiotic potential. Therefore, the present study was undertaken to isolate LAB from the phylloplanes of selected edible and culinary leaves and to evaluate their probiotic characteristics through a range of functional and physiological assays. The findings aim to contribute to the identification of novel LAB strains that may be used in the development of functional foods, dietary supplements, or natural bio preservatives.

## 2. Materials and methods

### 2.1. Plant Source

The different leaves *viz, chekkurmanis* (*Sauropus androgynus*), *betel* vine *(Piper betle*), *brahmi* (*Centella asiatica*), jackfruit (*Artocarpus heterophyllus*), *pandan* (*Pandanus amaryllifolius*), banana (*Musa paradisiaca*), turmeric (*Curcuma longa*) and castor (*Ricinus communis*) were collected from horticulture medicinal garden and around UAS, GKVK campus for isolation and enumeration of microorganisms.

### 2.2. Isolation of Microorganisms from the Phylloplanes of Leaf Samples

The standard plate count method (for small phylloplane samples *i*.*e*., *chekkurmanis, betel* vine, *brahmi*) and cotton swab technique (for large phylloplane samples *i*.*e*., *pandan*, jackfruit, banana, turmeric and castor) were employed for isolation of microorganisms. The leaves samples (*chekkurmanis, betel* vine, *brahmi*) were weighed one gram and transferred to sterile water blank. A sterile cotton swab was employed to sample the microflora from the surface of leaves within designated 10 cm^2^ areas. This semi-quantitative approach enables enumeration of the microorganisms per cm^2^ (Das, 2025; Jansson *et al*., 2020). The media *viz*, nutrient agar (NA) for bacteria, MRS agar for lactic acid bacteria and Martin’s Rose Bengal Agar (MRBA) for fungi were used for isolation of microorganisms (Bradbury, 1970; De Man *et al*., 1960; Waksman, 1922). The plates were incubated in an incubator at 25 °C for 48, 72 and 96 h for bacteria, LA bacteria and fungi respectively. The single discrete colonies were picked up and re-streaked on petri plates until pure cultures were obtained and were maintained on slants which were sub-cultured once in three months.

### 2.3. Probiotic parameters analysis

#### Bile Salt Tolerance

The tolerance of lactic acid bacterial (LAB) isolates to bile salts was assessed following the method of Ali *et al*. (2020) with minor modifications (Ali *et al*., 2020; Das, Banerjee, *et al*., 2025). Overnight cultures of LAB isolates grown in de Man, Rogosa and Sharpe (MRS) broth at 37□°C were inoculated (10 ^8^ cfu/mL) into fresh MRS broth supplemented with varying concentrations of bile salts (0.5, 1.0, 1.5, and 2.0%, w/v). Cultures were incubated at 37□°C for 24□h and 48 h. Uninoculated tubes served as controls. Growth was assessed by measuring optical density (OD) at 660□nm.

#### Acid Tolerance

Acid tolerance was evaluated according to Mulaw *et al*. (2019) (Mulaw *et al*., 2019). LAB isolates were cultured overnight in MRS broth and harvested by centrifugation (5000 rpm, 10 min, 4□°C). Pellets were washed twice with phosphate-buffered saline (PBS, pH 7.2) and resuspended in MRS broth adjusted to pH values of 2.0, 3.5, 5.0, and 7.0 using 1N HCl. Uninoculated tubes were used as controls, cultures were incubated at 37□°C, and growth was measured as OD at 660□nm at 24-hour intervals for 48 hours.

#### Temperature Tolerance

Temperature tolerance was determined following Ali *et al*. (2020) (Ali *et al*., 2020). Overnight cultures of LAB isolates were inoculated (10 ^8^ cfu/mL) into MRS broth and incubated at 25, 30, 37 and 40□°C for 24 and 48 hours. Uninoculated tubes were used as controls. Growth was quantified by OD measurement at 660□nm.

#### Haemolytic Activity

Haemolytic activity was tested using the method described by Bergey (1930) (Trujillo *et al*., 2015). LAB isolates were inoculated into wells (5□mm diameter) of sheep blood agar plates prepared using 5% (v/v) defibrinated sheep blood in Tryptic Soy Agar (TSA). Plates were incubated at 30□°C for 72 hours. Haemolysis was assessed by observing the diameter of the clear zone around the wells. Sterile distilled water served as a control.

#### Antibiotic Susceptibility

Antibiotic susceptibility was evaluated using the disc diffusion method described by Bauer *et al*. (1966) (Bauer *et al*., 1966). LAB cultures were grown in MRS broth for four days at 37□°C and adjusted to a final population of 5 × 10□ cfu/mL. MRS agar was seeded with 5mL of culture per 100mL medium. Antibiotic discs (ampicillin 10□μg, azithromycin 15□μg, chloramphenicol 30□μg, gentamycin 10□μg, nystatin 50μg, and amphotericin-B 20μg) were placed on agar surfaces and incubated at 37□°C for 48 hours. Zone of inhibition were measured in centimetres.

#### NaCl Tolerance

NaCl tolerance was tested as described by Escamilla-Montes *et al*. (2015) (Escamilla-Montes *et al*., 2015). LAB isolates were inoculated (10^8^ cfu/mL) into MRS broth supplemented with 5, 6, and 7% (w/v) NaCl and incubated at 37□°C for 24 and 48 hours. Growth was measured spectrophotometrically at 600□nm. Tubes without NaCl served as controls.

#### Phenol Resistance

Phenol resistance was assessed following the protocol of Jena *et al*. (2013) (Jena *et al*., 2013). LAB isolates were grown in MRS broth containing 0.4% (v/v) phenol and incubated at 37□°C for 24 hours. Cell viability was determined by standard plate count method and expressed as log CFU/mL.

#### Auto-aggregation ability

Auto-aggregation was evaluated according to Zommiti *et al*. (2017) (Zommiti *et al*., 2017). Overnight cultures were centrifuged (8000 rpm, 10 min, 4□°C), washed twice in PBS, and resuspended in the same buffer. Absorbance of the upper suspension was recorded at 600□nm at 0, 1, 2, 3, 4, and 5 hours. Auto-aggregation (%) was calculated as:

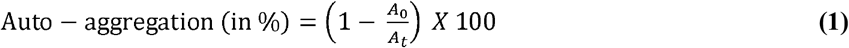

Where *A*□ is the absorbance at 0 h and *A*□ at a specific time.

#### Cell Surface Hydrophobicity

Cell surface hydrophobicity was determined using microbial adhesion to hydrocarbons (MATH) method (Rokana *et al*., 2018) (Rokana *et al*., 2018). LAB cells were harvested, washed with PBS, and absorbance (A□) recorded at 600□nm. A 3mL cell suspension was mixed with 1mL xylene and incubated at 37□°C for 1 hour without agitation. After phase separation, aqueous phase absorbance (A□) was measured. Hydrophobicity (%) was calculated as:

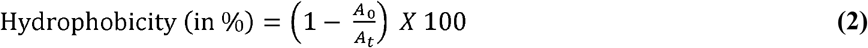

### 2.4. Statistical Analysis

Data were analysed using analysis of variance (ANOVA) with OPSTAT 2.0 software. Differences among means were assessed using Duncan’s Multiple Range Test at a significance level of *p* ≤ 0.05 (Duncan, 1955).

## 3. Results and discussion

LAB isolated from phylloplanes that exhibited significant antibacterial and antifungal activity were selected and labelled as LAB-1 to LAB-15 for further evaluation of their probiotic potential. To be considered probiotic, LAB must remain viable and functional under the harsh conditions of the gastrointestinal tract (GIT). Key attributes assessed included haemolytic activity, antibiotic sensitivity, surface hydrophobicity, auto-aggregation, NaCl and phenol tolerance, temperature and acid tolerance, and bile salt resistance.

### 3.1. Haemolytic Activity

LAB must be non-haemolytic to be considered safe for probiotic use. Haemolysis was assessed on trypticase soy agar supplemented with 5% sheep blood after four days of incubation. Haemolytic activity is categorized as α-haemolysis (partial, greenish zone), β-haemolysis (complete, clear zone), or γ-haemolysis (none). All tested LAB isolates showed γ-haemolysis, indicating no haemolytic activity (**Table 1**). This is consistent with findings by Tatsaporn and Kornkanok (2020), and Chandel (2019), who reported negative haemolytic activity for *Pediococcus pentosaceus, Enterococcus faecium*, and *Lactobacillus* spp., confirming their safety for probiotic application (Chandel *et al*., 2019; Tatsaporn & Kornkanok, 2020).

**Table 1:**
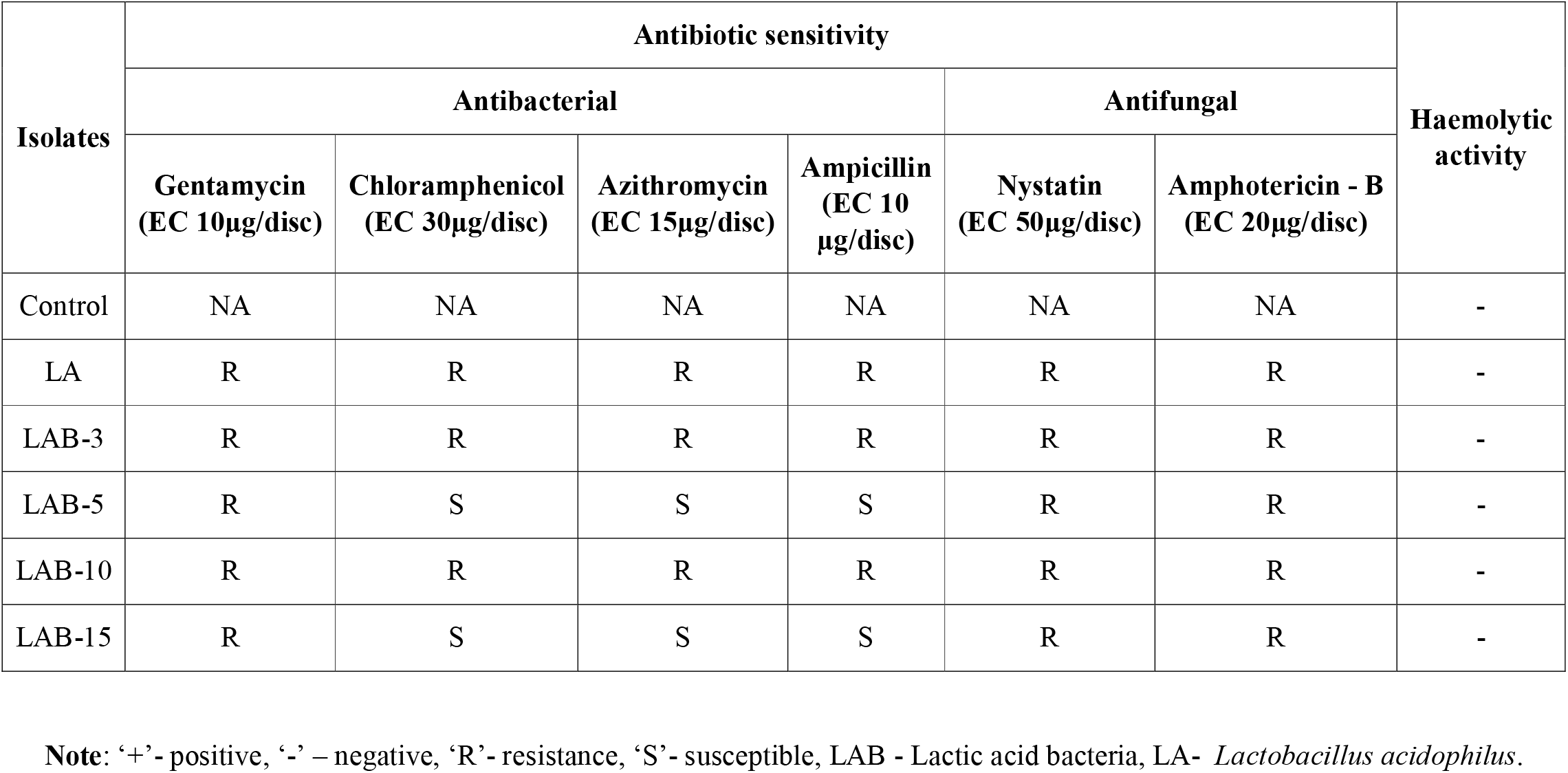
Antibiotic sensitivity and haemolytic activity of LA bacterial isolates.

### 3.2. Antibiotic Sensitivity

Isolates *Lactobacillus acidophilus*, LAB-3, and LAB-10 were resistant to six antibiotics—gentamicin, chloramphenicol, azithromycin, ampicillin, nystatin, and amphotericin B (**Table 1**). All isolates were resistant to the two antifungal agents tested. Dowarah *et al*. (2018) reported similar findings in LAB isolated from piglet feces, with resistance to ciprofloxacin, ofloxacin, gatifloxacin, vancomycin, and co-trimoxazole, but sensitivity to penicillin and other antibiotics. Likewise, Sirichoat *et al*. (2020) reported acquired resistance among vaginal LAB strains (Sirichoat *et al*., 2020).

### 3.3. Cell Surface Hydrophobicity

Hydrophobicity is an important factor in bacterial adhesion to intestinal epithelial cells. The isolates showed hydrophobicity ranging from 19% to 72% (**Table 2**). *Lactobacillus acidophilus* showed the highest hydrophobicity (72%), followed by LAB-10 (60%). LAB-3 and LAB-5 had less than 40%. Dowarah *et al*. (2018) reported 64% hydrophobicity in *Lacp28* (Dowarah *et al*., 2018), while Meena *et al*. (2022) observed 75.3 ± 2.76% in *Lactobacillus delbrueckii subsp. bulgaricus* KMUDR1 (Meena *et al*., 2022).

**Table 2:** Percent auto-aggregation and cell hydrophobicity of LA bacterial cultures at intervals.

| Isolates | Auto aggregation (%) |  |  |  |  | Cell hydrophobicity (%) |
| --- | --- | --- | --- | --- | --- | --- |
|  | 1h | 2h | 3h | 4h | 5h | 24h |
| Control | 00 | 00 | 00 | 00 | 00 | 00 |
| LA | 19 | 24 | 30 | 31 | 42 | 72 |
| LAB-3 | 2 | 4 | 4 | 7 | 11 | 38 |
| LAB-5 | 10 | 12 | 19 | 24 | 28 | 19 |
| LAB-10 | 10 | 15 | 25 | 29 | 36 | 60 |
| LAB-15 | 19 | 21 | 22 | 24 | 32 | 57 |

**Note:** LAB- Lactic acid bacteria, numerical values are mean of three replicates, LA- *Lactobacillus acidophilus*

### 3.4. Auto-Aggregation

Auto-aggregation correlates with bacterial adherence capability. After 5 hours, auto-aggregation ranged from 11–42% (**Table 2**). *Lactobacillus acidophilus* exhibited the highest (42%), followed by LAB-10 (36%). Xu *et al*. (2009) observed 51.8% auto-aggregation in *Bifidobacterium longum* B6. Topcu *et al*. (2020) found 17.62% auto-aggregation in *Pediococcus pentosaceus* K41 (Topçu *et al*., 2020).

### 3.5. NaCl Tolerance

NaCl tolerance is essential for LAB survival in osmotic stress conditions. All isolates grew at 4–6% NaCl, with growth declining at higher concentrations due to salt’s inhibitory effects (**Table 3**). *Lactobacillus acidophilus* showed the highest growth, followed closely by LAB-10. Pundir *et al*. (2013) similarly reported LAB growth up to 6–6.5% NaCl and reduced growth at 7%, with no survival at 8–10% (Ram Kumar Pundir, 2013).

**Table 3:** Absorbance of LA bacterial cultures at different NaCl and phenol concentrations.

| Isolates |  |  | Control | LA | LAB- 3 | LAB- 5 | LAB- 10 | LAB- 15 |
| --- | --- | --- | --- | --- | --- | --- | --- | --- |
| NaCl concentrations (%) | 4 | 3 h | 0.00<sup>d</sup> | 0.73<sup>a</sup> | 0.18<sup>c</sup> | 0.15<sup>c</sup> | 0.58<sup>b</sup> | 0.53<sup>b</sup> |
|  |  | 6 h | 0.00<sup>d</sup> | 0.87<sup>a</sup> | 0.21<sup>c</sup> | 0.17<sup>c</sup> | 0.79<sup>b</sup> | 0.77<sup>b</sup> |
|  |  | 24 h | 0.00<sup>d</sup> | 0.89<sup>a</sup> | 0.21<sup>c</sup> | 0.17<sup>c</sup> | 0.81<sup>b</sup> | 0.79<sup>b</sup> |
|  | 5 | 3 h | 0.00<sup>e</sup> | 0.7<sup>a</sup> | 0.15<sup>d</sup> | 0.14<sup>d</sup> | 0.55<sup>b</sup> | 0.48<sup>c</sup> |
|  |  | 6 h | 0.00<sup>d</sup> | 0.86<sup>a</sup> | 0.20<sup>c</sup> | 0.15<sup>c</sup> | 0.76<sup>b</sup> | 0.72<sup>b</sup> |
|  |  | 24 h | 0.00<sup>d</sup> | 0.88<sup>a</sup> | 0.20<sup>c</sup> | 0.15<sup>c</sup> | 0.77<sup>b</sup> | 0.74<sup>b</sup> |
|  | 6 | 3 h | 0.00<sup>e</sup> | 0.68<sup>a</sup> | 0.15<sup>d</sup> | 0.13<sup>d</sup> | 0.53<sup>b</sup> | 0.47<sup>c</sup> |
|  |  | 6 h | 0.00<sup>d</sup> | 0.83<sup>a</sup> | 0.16<sup>c</sup> | 0.15<sup>c</sup> | 0.73<sup>b</sup> | 0.71<sup>b</sup> |
|  |  | 24 h | 0.00<sup>d</sup> | 0.85<sup>a</sup> | 0.16<sup>c</sup> | 0.15<sup>c</sup> | 0.75<sup>b</sup> | 0.72<sup>b</sup> |
| Phenol (0.4 %) |  | 24 h | 0.00<sup>b</sup> | 0.10<sup>a</sup> | 0.07<sup>a</sup> | 0.07<sup>a</sup> | 0.08<sup>a</sup> | 0.08<sup>a</sup> |
|  |  | 0 h | 0.00<sup>e</sup> | 0.31<sup>c</sup> | 0.27<sup>c</sup> | 0.14<sup>d</sup> | 0.50<sup>a</sup> | 0.44<sup>b</sup> |

**Note:** LAB- Lactic acid bacteria, LA- *Lactobacillus acidophilus*, numerical values are mean of three replicates. The treatment means with the different letters as superscript in the same columns as determined by DMRT ( $p \leq 0.05$ ).

### 3.6. Phenol Tolerance

LAB must resist toxic phenolic compounds produced during digestion. Phenol tolerance was evaluated at 0.4% phenol after 24 hours at 600 nm. LAB-10 exhibited the highest tolerance (0.50), followed by LAB-15 (0.44) (**Table 3**). Vizoso-Pinto *et al*. (2006) and Reuben *et al*. (2019) also reported phenol resistance among LAB strains, with varying OD values indicating survivability.

### 3.7. Temperature Tolerance

Survivability at body temperature is a key probiotic trait. *Lactobacillus acidophilus*, LAB-10, and LAB-15 showed optimal growth at 37–40 °C, while LAB-3 and LAB-5 had suboptimal growth at all tested temperatures (**Figure 1**). Reuben *et al*. (2019) and Divyashree *et al*. (2021) reported similar findings, with optimal LAB growth at 37 °C after 24 hours (Divyashree *et al*., 2021; Reuben *et al*., 2019).

**Figure 1:**
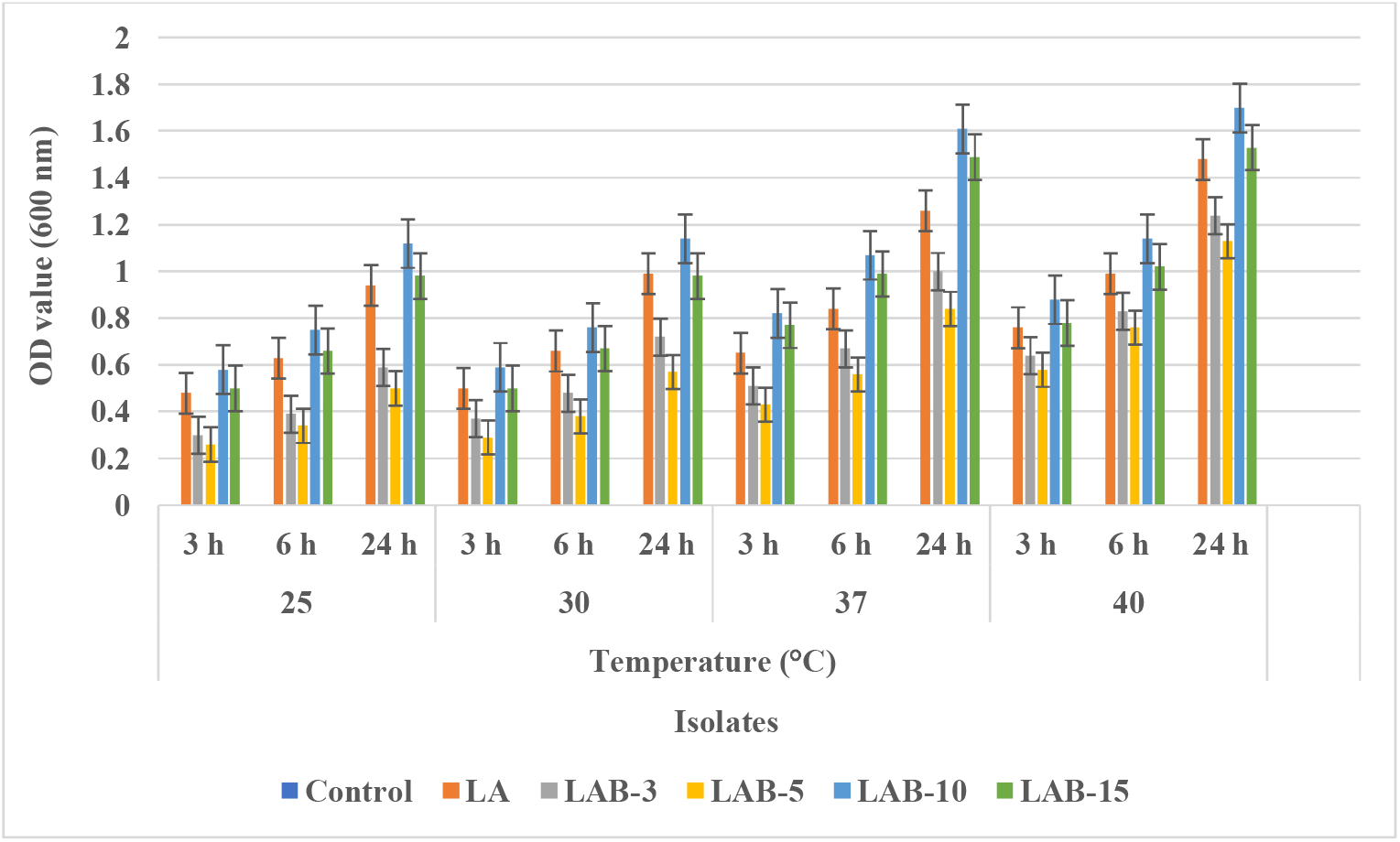
Effect of temperature on growth of LA bacterial isolates in MRS broth at intervals.

### 3.8. Acid Tolerance

Acid tolerance enables LAB to survive gastric conditions. All isolates survived at pH values ranging from 2.0 to 8.0, with better survival at pH 2.0–3.5. LAB-10 showed the highest survival rate, followed by *Lactobacillus acidophilus*. LAB-5 had the lowest viability across all pH levels (**Figure 2**). Comparable results were reported by Chen *et al*. (2020) and Vanniyasingam *et al*. (2019), confirming the acid tolerance of *Lactobacillus plantarum* and other LAB strains at pH as low as 2.0 (Chen *et al*., 2020; Vanniyasingam *et al*., 2019).

**Figure 2:**
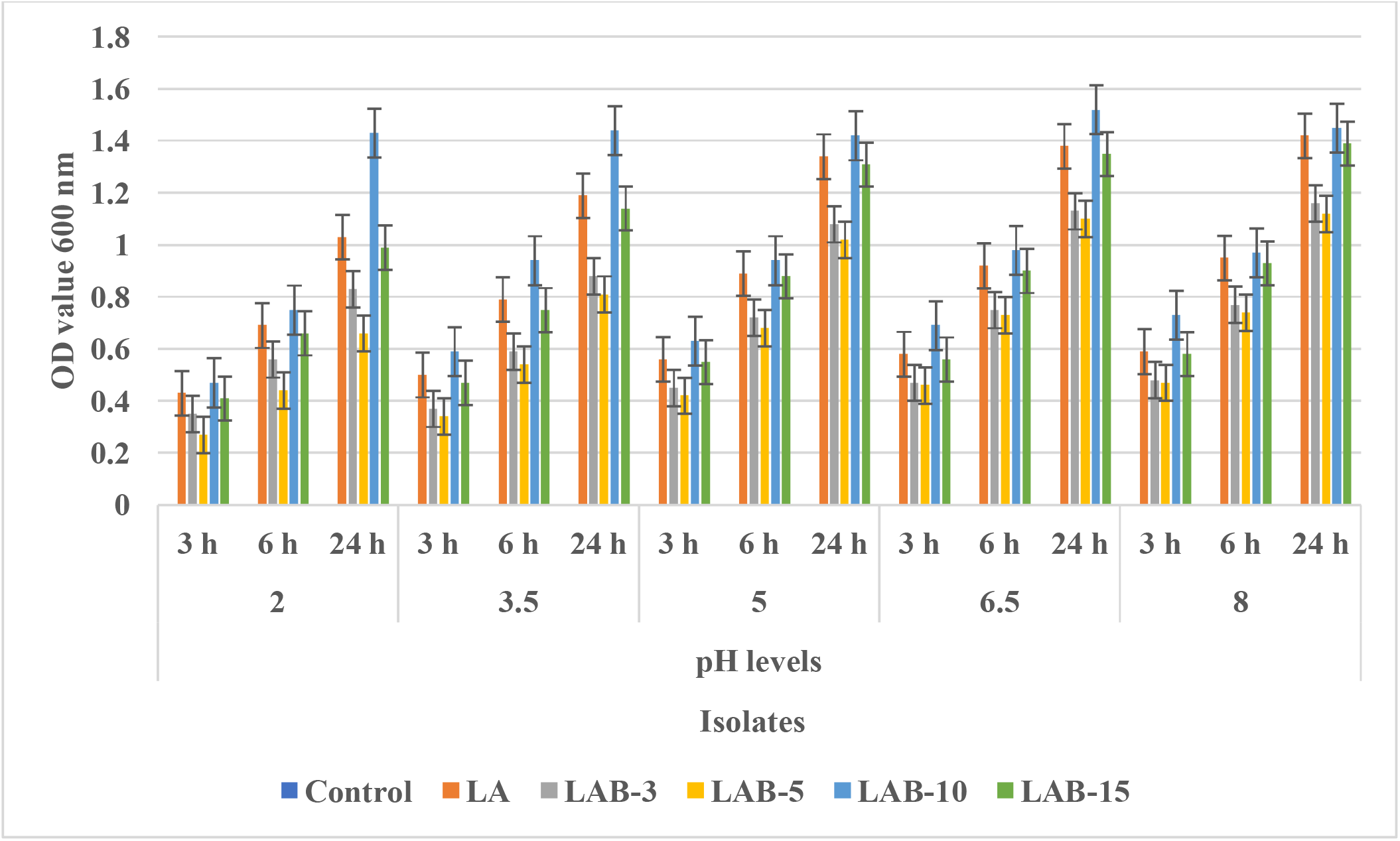
Effect of pH on growth of LA bacterial isolates in MRS broth at intervals.

### 3.9. Bile Salt Tolerance

Bile tolerance is necessary for colonization and metabolic activity in the intestine. The isolates were tested at 0.5–2.0% bile salt concentrations over 3, 6, and 24 hours. All isolates were bile-tolerant, with LAB-10 exhibiting the highest tolerance, followed by *Lactobacillus acidophilus*. LAB-5 showed the lowest (**Figure 3**). Similar bile resistance was reported by Chen *et al*. (2020) and Menconi *et al*. (2014), who noted viability reduction with increasing bile concentrations, but confirmed overall tolerance of the strains tested (Chen *et al*., 2020; Menconi *et al*., 2014).

**Figure 3:**
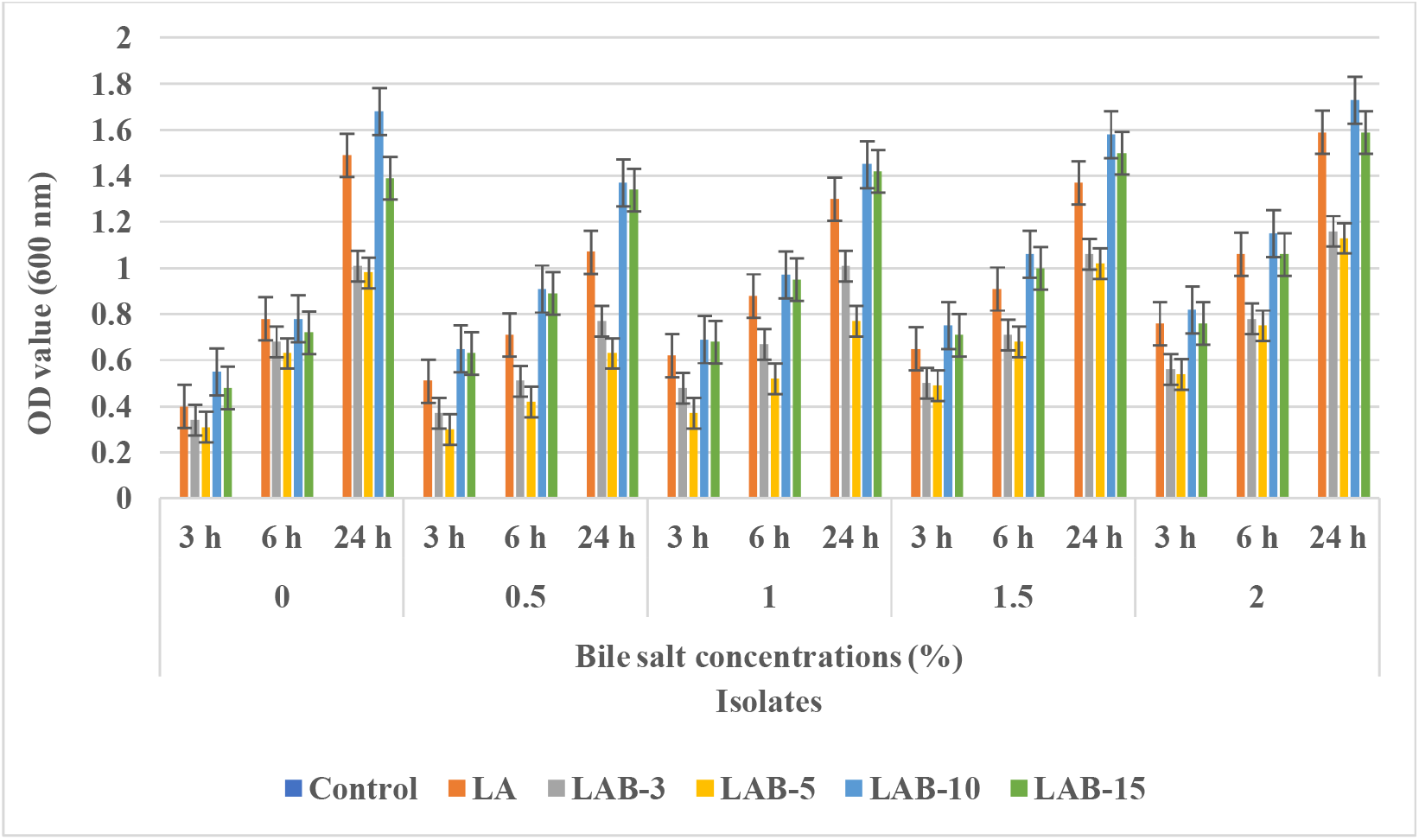
Effect of bile salt concentrations on growth of LA bacterial isolates in MRS broth at intervals.

## 4. Conclusion

The present study demonstrates that lactic acid bacteria (LAB) isolated from the phylloplanes of raw edible and culinary leaves possess promising probiotic attributes. Selected LAB isolates that initially exhibited notable antibacterial and antifungal activity were further evaluated for their tolerance to gastrointestinal stress factors and their potential health-promoting properties. All isolates were non-haemolytic, confirming their safety and exhibited varying degrees of resistance to commonly used antibiotics and antifungal agents. Among them, *Lactobacillus acidophilus* and isolate LAB-10 showed superior probiotic potential by demonstrating high cell surface hydrophobicity, significant auto-aggregation ability, and strong tolerance to NaCl, phenol, bile salts, and acidic pH conditions. These isolates also exhibited optimal growth at temperatures simulating the human body. The ability of these strains to survive under gastrointestinal-like conditions highlights their potential as effective probiotic candidates. Further *in vivo* studies and functional assessments are warranted to confirm their application in food and health industries.

## Declaration

We confirm that the plant materials used in our study do **not** include any species listed as endangered or protected under the IUCN Red List or the Convention on the Trade in Endangered Species of Wild Fauna and Flora (CITES). Accordingly, no special permissions or ethical clearances were required under these guidelines.

## Data Availability

The authors confirm that the data supporting the findings of this study are available within the article. Raw data supporting the findings of this study are available from the corresponding author upon reasonable request.

## Acknowledgement

None

## Declaration of generative AI and AI-assisted technologies in the writing process

The writing of this research paper involved the use of generative AI and AI-assisted technologies only to enhance the clarity, coherence, and overall quality of the manuscript. The authors acknowledge the contributions of AI in the writing process only. All interpretations and conclusions drawn in this manuscript are the sole responsibility of the author.

## Ethics Statement

This research did not involve any human participants or animal subjects, and therefore, no ethical approval was required.

## Author Contributions

***Shashank S***: Conceptualization, Methodology, Formal Analysis, Writing—Original Draft, Writing—Review & Editing. ***Uday Kumar B V***: Writing—Review & Editing. ***Uddalak Das***: Project administration. All the authors have read and agreed to the published version of the manuscript.

